# Metagenomic classification with KrakenUniq on low-memory computers

**DOI:** 10.1101/2022.06.01.494344

**Authors:** Christopher Pockrandt, Aleksey V. Zimin, Steven L. Salzberg

## Abstract

Kraken and KrakenUniq are widely-used tools for classifying metagenomics sequences. A key requirement for these systems is a database containing all *k*-mers from all genomes that the users want to be able to detect, where *k* = 31 by default. This database can be very large, easily exceeding 100 gigabytes (GB) and sometimes 400 GB. Previously, Kraken and KrakenUniq required loading the entire database into main memory (RAM), and if RAM was insufficient, they used memory mapping, which significantly increased the running time for large datasets. We have implemented a new algorithm in KrakenUniq that allows it to load and process the database in chunks, with only a modest increase in running time. This enhancement now makes it feasible to run KrakenUniq on very large datasets and huge databases on virtually any computer, even a laptop, while providing the same very high classification accuracy as the previous system.

---

The GenBank genome repository currently contains over 400,000 prokaryotic genomes and over 20,000 eukaryotes, including thousands of microbial eukaryotes such as fungi and protists. To take advantage of this ever-growing variety of microbial sequences, metagenomic sequence analysis methods must create customized databases that capture all of this sequence diversity. Tools such as Kraken [WS14] and KrakenUniq [BBS18] classify DNA or RNA sequencing reads against a pre-built database of genomes using an exact *k*-mer matching strategy that is not only highly accurate but that, because it avoids the step of sequence alignment, makes both systems extremely fast.

The Kraken database is a customized, compressed data structure that associates a unique taxonomy identifier with every single *k*-mer in every genome in the database. (Note that both Kraken and KrakenUniq use the same database design, so we will refer to both as Kraken databases.) If a *k*-mer occurs in two or more genomes, then the database stores the taxonomy ID associated with the lowest common ancestor of those genomes. This strategy means that only a single ID is attached to each *k*-mer.

However, with the number of genomes available today, a standard Kraken database will contain billions of *k*-mers, and even with careful compression this data structure can grow very large. A key requirement for the speed of the Kraken algorithm (which is 900 times faster than MegaBlast [WS14]) is the loading of the entire database into main memory. For the large databases and read datasets that are commonly used in metagenomics experiments today, this requires dedicated machines with large amounts of RAM (e.g., exceeding 100 GB or even 400 GB), without which classification becomes slow and impractical. The newer Kraken2 system [WLL19] achieves a significantly lower memory footprint by using probabilistic data structures to reduce the database size, at the cost of slightly lower accuracy than KrakenUniq. This reduction in accuracy includes a very small but non-zero false positive rate (i.e., where the system incorrectly reports that a *k*-mer is present in a particular genome), which is problematic for certain applications that require very high precision. In particular, when metagenomic sequencing is used for the diagnosis of infections in a clinical setting [SBK^+^16], the pathogen of interest might be detected from just a handful of reads. In that scenario, even a few false positives can be confusing, and KrakenUniq is the preferred method rather than Kraken2.

By default, KrakenUniq performs memory mapping to load the database; i.e., it does not load the entire database into main memory. (Kraken 1 employs the same strategy.) This makes classification of larger read datasets much slower, but it allows KrakenUniq to run on machines with low available main memory. If enough free RAM is available to hold the entire database in main memory, users are recommended to explicitly load the entire database prior to classification using the flag `--preload`, which dramatically speeds up the classification (see **Table 1**).

**Table 1:** Running times for classifying 9.4 million reads (from a human eye metagenome, SRR12486990) with 8 threads using KrakenUniq in different modes. The database size was 392 GB, and it consisted of all complete bacterial, archeal, and viral genomes in RefSeq from 2020, 46 selected eukaryotic human pathogens [LS18]), as well the human genome and a collection of common vector sequences. In the database chunking experiments (using –preload-size) KrakenUniq loaded the database into main memory in 49, 25, 13 and 7 chunks (respectively).

| Mode | Running time |
| --- | --- |
| default (memory mapping) | 48 hours |
| `--preload` | 14 min |
| `--preload-size 8G` | 47 min |
| `--preload-size 16G` | 32 min |
| `--preload-size 32G` | 25 min |
| `--preload-size 64G` | 19 min |

To improve KrakenUniq’s performance when not enough main memory is available to load the entire database into RAM, we have added a new capability to KrakenUniq, which we call database chunking. This new feature is released as KrakenUniq v0.7 at: https://github.com/fbreitwieser/krakenuniq (Github) https://anaconda.org/bioconda/krakenuniq (Conda)

## Database chunking

Under this new algorithm, KrakenUniq only loads a chunk of the database into memory at a time, based on the amount of available memory. It then iterates over all of the reads provided as input and looks up all *k*-mers in those reads that are matching in this database chunk. This process is repeated until the entire database has been processed. The k-mer lookups are then merged, and reads are classified based on the results of the full database. Classification results will be identical to running in the default mode; i.e., database chunking does not alter the output.

This new feature makes it feasible to run KrakenUniq on very large datasets and huge databases on virtually any computer, even a laptop, while providing exact classifications that are identical to those of KrakenUniq in its other modes. Users can employ this feature by using `--preload-size` to specify the amount of available main memory that they want to use for loading chunks of the database, e.g., `--preload-size 8G` or `--preload-size 500M`.

Running times and speedups can vary significantly depending on the type of storage (e.g., databases on network storage can take longer to load) and the size of the read dataset (i.e., classifying a small number of reads does not justify preloading the entire database, especially not on slow storage). **Table 1** shows that in a typical use case, loading the entire database is far faster than memory mapping (14 minutes versus 48 hours). Loading the database by chunks adds overhead because of the need to iterate over the reads multiple times, but is still comparable to pre-loading the entire database and highly recommended when not enough main memory is available. For example, limiting the database to 8G, which means it can be loaded even on a standard laptop computer, increased the running time only about 3.4-fold, even though the database was broken into 49 chunks. The format of the databases used by the new algorithm has not changed, hence all previously built databases for Kraken and KrakenUniq can be used.

This feature has only been added to KrakenUniq, and not to Kraken, which is no longer actively maintained. Because KrakenUniq offers more features, shares the same implementation with Kraken and produces the same output, we highly recommend that users upgrade from Kraken to KrakenUniq.

## Acknowledgements

This work was supported in part by NIH grants R35-GM130151 and R01-HG006677.

## References

[BBS18] F.P. Breitwieser, D.N. Baker, and S.L. Salzberg. KrakenUniq: confident and fast metagenomics classification using unique k-mer counts. Genome Biology, 19(1):1–10, 2018.

[LS18] Jennifer Lu and Steven L. Salzberg. Removing contaminants from databases of draft genomes. PLoS Computational Biology, 14(6):e1006277, 2018.

[SBK+16] SL Salzberg, FP Breitwieser, A Kumar, H Hao, P Burger, FJ Rodriguez, M Lim, A Quiñones-Hinojosa, GL Gallia, JA Tornheim, MT Melia, CL Sears, and CA Pardo. Next-generation sequencing in neuropathologic diagnosis of infections of the nervous system. Neurology: Neuroimmunology and Neuroinflammation, 3(4):e251, 2016.

[WLL19] Derrick E. Wood, Jennifer Lu, and Ben Langmead. Improved metagenomic analysis with kraken 2. Genome Biology, 20(1):1–13, 2019.

[WS14] Derrick E. Wood and Steven L. Salzberg. Kraken: ultrafast metagenomic sequence classification using exact alignments. Genome Biology, 15(3):1–12, 2014.

